# Vitamin A-treated natural killer cells reduce interferon-gamma production and support regulatory T cell differentiation

**DOI:** 10.1101/2022.12.05.519129

**Authors:** Mingeum Jeong, Jia-Xiang See, Carolina De La Torre, Adelheid Cerwenka, Ana Stojanovic

## Abstract

Natural killer (NK) cells are innate cytotoxic lymphocytes that contribute to immune responses against stressed, transformed or infected cells. NK cell effector functions are regulated by microenvironmental factors, including cytokines, metabolites and nutrients. Vitamin A is an essential micronutrient that plays an indispensable role in embryogenesis and development, but was also reported to regulate immune responses. However, the role of vitamin A in regulating NK cell functions remains poorly understood. Here, we show that the most prevalent vitamin A metabolite, all-*trans* retinoic acid (atRA), induces transcriptional and functional changes in NK cells leading to altered metabolism and reduced IFN-γ production in response to a wide range of stimuli. atRA-exposed NK cells display a reduced ability to support dendritic cell (DC) maturation and to eliminate immature DCs. Moreover, they support the polarization and proliferation of regulatory T cells. These results imply that in vitamin A-enriched environments, NK cells can acquire regulatory-like functions and might promote tolerogenic immunity and/or immunosuppression.

## Introduction

NK cells are innate lymphocytes that play important roles in anti-tumor and anti-viral immunity [1]. The balance of signals delivered by the surface activating and inhibitory receptors determines NK cell activation [2]. NK cells can eliminate target cells via receptor-triggered secretion of cytotoxic granules or by inducing apoptosis through the engagement of death receptors expressed by target cells [3]. Furthermore, soluble factors can modulate NK cell functions [4]. For example, cytokines IL-12 and IL-18 synergistically trigger NK cell activation and production of pro-inflammatory cytokine IFN-γ [5].

NK cells contribute to regulation of adaptive immunity through direct interaction with T cells, as well as by regulating the functions of dendritic cells (DCs) [6]. IL-12 and IL-18 released by DCs, or the engagement of the activating receptor NKp30 by its ligand(s) expressed by DCs, were shown to enhance NK cell cytotoxicity and cytokine production, including IFN-γ and TNF-α [7].Reciprocally, cytokines secreted by NK cells induce DC maturation, characterized by elevated expression of MHC class I and co-stimulatory molecules, while at a high NK/DC ratio, NK cells were shown to kill immature DCs [8–10]. During T cell priming, NK cells are recruited to lymph nodes and support the differentiation of Th1 cells by providing IFN-γ [11, 12]. On the contrary, NK cells can restrict the responses of activated T cells via NKG2D-mediated cytotoxicity [13, 14].

Microenvironmental factors, including micronutrients or oxygen tension, can modulate activation and effector functions of NK cells [15, 16]. Vitamin A is an essential fat-soluble micronutrient that is digested and absorbed in the small intestine, and stored mainly in the liver [17]. NK cells are recruited to these tissues upon inflammation [18–20], and therefore, can be exposed to vitamin A-enriched microenvironment. Previous studies have demonstrated that animals fed with vitamin A-deficient diet showed a decreased number of NK cells in blood, while splenic NK cells displayed impaired effector functions, which could be restored by the oral administration of retinoic acid [21, 22]. Contrarily, the cytotoxicity of the human NK92 cell line was suppressed by *in vitro* treatment with atRA [23]. Thus, the exact roles of vitamin A and its metabolites in regulating NK cell phenotype and functions remains incompletely understood.

Here, we show that NK cells treated with the most prevalent vitamin A metabolite, all-*trans* retinoic acid (atRA), acquire regulatory-like functions. atRA-treated NK cells produce less IFN-γ, favor an immature DC phenotype, and support the differentiation of regulatory T (Treg) and regulatory Th17 cells, as well as their proliferation. Our data provide a novel insight in NK cell regulation by vitamin A, which might affect not only tissue homeostasis and immune tolerance, but also support immunosuppression during an active immune response.

## Results

### atRA induces transcriptional and phenotypic changes in NK cells

To address the effect of vitamin A, mouse splenic NK cells exposed for 7 days to the active form of vitamin A metabolite, atRA, or the equivalent volume of solvent, DMSO, were subjected to transcriptome analysis. atRA-treated NK displayed over 400 differentially regulated transcripts (Fig. 1A). Expression of genes involved in retinoic acid (RA) and lipid metabolism (Dhr3, Dgat1, and Pparg) was upregulated, and the expression of genes involved in NK cell activation and effector functions (Ifng, Il18r1, Tnf, Klrb1c, and Ncr1) was downregulated upon the exposure to atRA (Fig. 1A-B). Accordingly, retinol metabolism pathway was enriched, while the NFκB and NK cell-mediated cytotoxicity pathways were downregulated in atRA-treated NK cells compared to controls (Fig. 1C).

**Figure 1.**
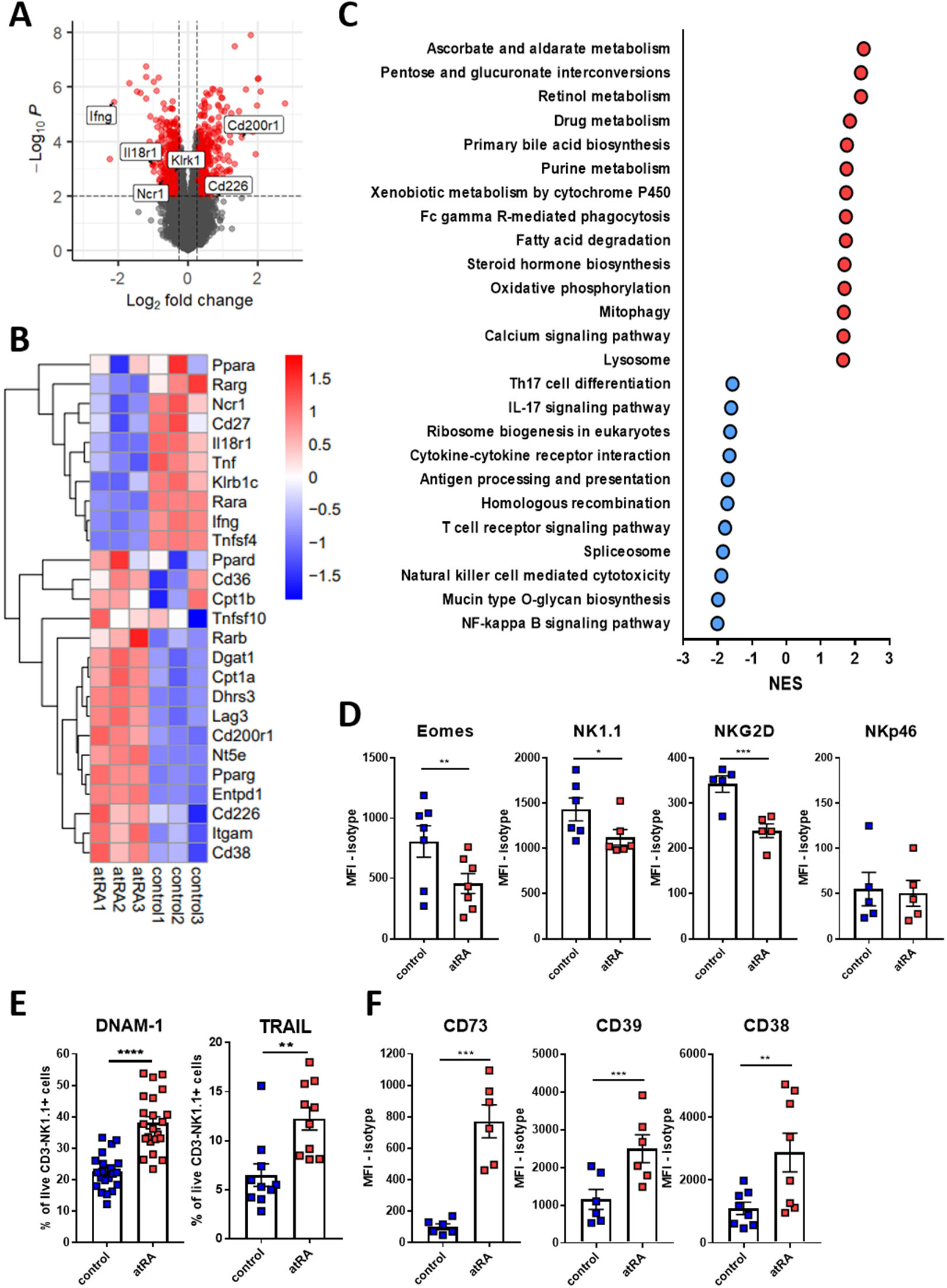
atRA induces transcriptomic and phenotypic alterations in NK cells. **(A)** NK cells were cultured with 1 µM of atRA (or DMSO as a control solvent) in the presence of IL-2 for 7 days. The differential transcript expression in atRA-treated compared to control NK cells. Cut-off (red): Fold-Change > 1.2 or < −1.2, and p-value<0.01. **(B)** Hierarchical clustering of selected differentially expressed candidate transcripts. **(C)** Pathway enrichment analysis based on Kyoto Encyclopedia of Genes and Genomes (KEGG) for atRA-treated compared to control NK cells (FC > 1.5 or < 1.5 and adjusted p-value < 0.05). NES, normalized enrichment score; MFI, mean florescence intensity. **(D-F)** Control and atRA-treated NK cells were analyzed by flow cytometry (n=5-22). Graphs indicate mean ± SEM. *, p<0.05; **, p<0.01; ***, p<0.001; ****, p<0.0001 (paired Student’s t-test; Welch’s t-test for analysis of TRAIL expression).

Next, we analyzed protein expression of selected molecules whose transcripts were differentially expressed in atRA-treated NK cells. The expression of the transcription factor Eomes was decreased in atRA-treated NK cells compared to controls, as well as the expression of activating receptors NK1.1 and NKG2D, while the expression of NKp46 remained unchanged (Fig. 1D). atRA-treated NK cells comprised higher frequencies of DNAM-1- and TRAIL-expressing cells (Fig. 1E). Similarly, the expression of nucleotide-metabolizing ectoenzymes CD73, CD39, and CD38 increased over time during culture of NK cells with atRA (Fig. 1F and Fig. S1). As RA was reported to induce the expression of gut-homing receptors in T cells and group 3 innate lymphoid cells (ILC3s) [24, 25], we examined the expression of adhesion molecules and homing-receptors on atRA-treated NK cells. Among atRA-treated NK cells, CXCR3-expressing cells were enriched, CD62L- and CCR9-expressing cells were reduced, while the frequencies of CXCR6-expressing cells remained comparable to control-treated NK cells (Fig. S2). Together, these data demonstrate that the exposure to atRA induces comprehensive transcriptional and phenotypic alterations in NK cells.

### atRA-treated NK cells display altered metabolism

Mitochondria contribute to cellular metabolism by oxidizing pyruvate, fatty acids, and amino acids to generate energy [26]. As atRA-treated NK cells displayed altered expression of genes involved in several metabolic pathways (Fig. 1C), we assessed the features of mitochondria in NK cells upon treatment with atRA. We detected a reduced mitochondrial mass and membrane potential, as well as reduced production of reactive oxygen species by atRA-treated NK cells (Fig. 2A). Accordingly, mitochondrial respiration and glycolysis, measured via the oxygen consumption rate (OCR) and the extracellular acidification rate (ECAR), respectively, were altered; atRA-treated NK cells showed reduced basal and maximal OCR, and reduced ECAR (Fig. 2B-C). Furthermore, atRA-treated NK cells were smaller (Fig. 2D), while their expansion rate in culture (Fig. S3A) and the uptake of fatty acids, remained unchanged compared to controls (Fig. S3B). These results indicate that the exposure to atRA resulted reduced mitochondrial mass and function, as well as altered metabolism of NK cells.

**Figure 2.**
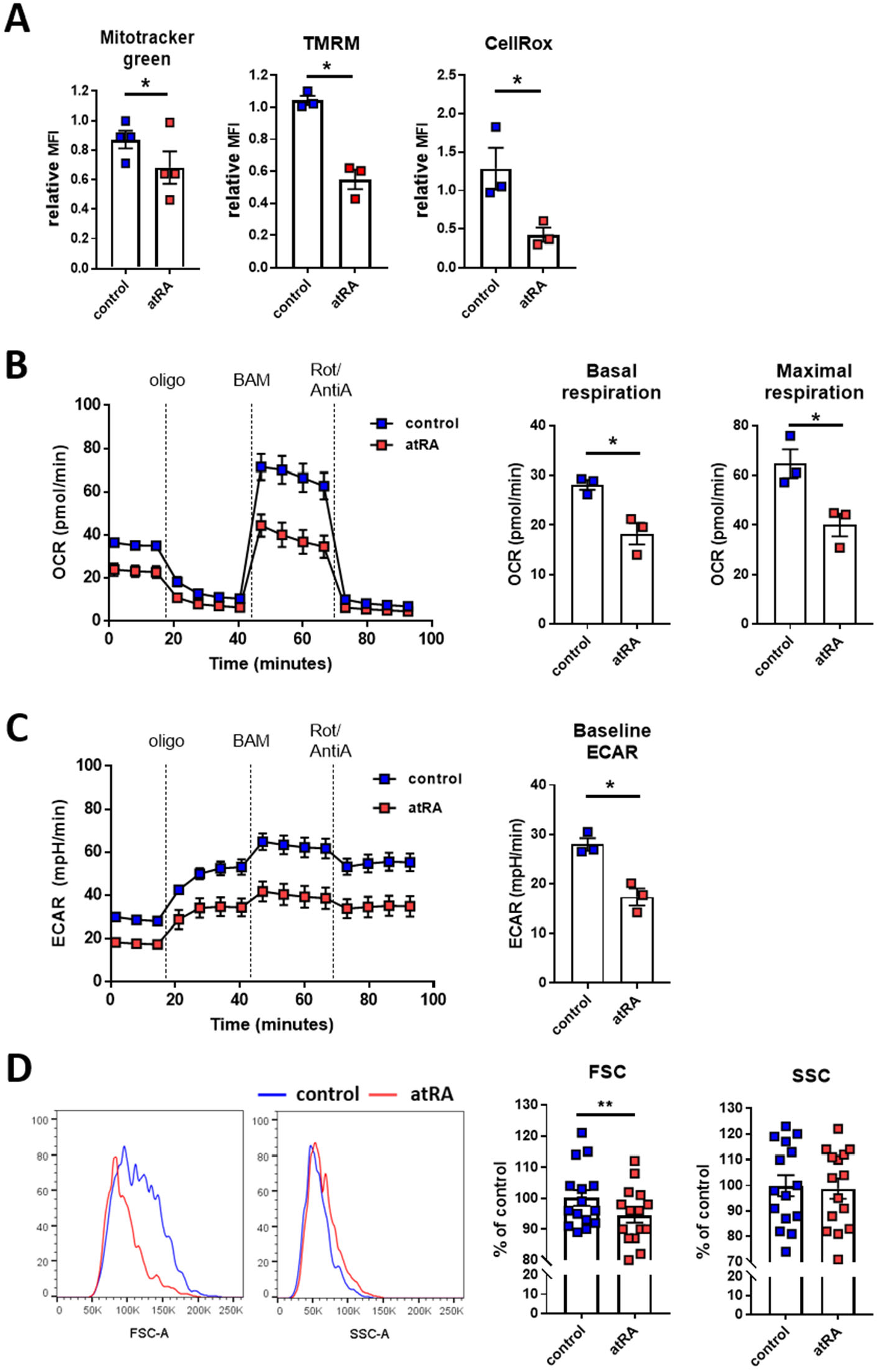
atRA-treated NK cells show differential mitochondrial phenotype and reduced metabolic activity. **(A)** NK cells were cultured with 1 µM of atRA (or DMSO as a control solvent) in the presence of IL-2 for 7 days. Cells were analyzed for mitochondrial mass (Mitotracker™), mitochondrial membrane potential (TMRM) and reactive oxygen species (ROS) production (CellRox™) by flow cytometry. (n=3-4; mean ± SEM; *p<0.05by paired Student’s t-test) **(B-C)** Oxygen consumption rate (OCR) and extracellular acidification rate (ECAR) were analyzed upon treatment with oligomycin (oligo), BAM15, and Rotenone and Antimycin A (Rot/AntiA). Graphs show OCR and ECAR over 100 min time-lapse, and quantification of basal and maximal respiration, and baseline ECAR (n=3, mean ± SEM; *p<0.05 by paired Student’s t-test). **(D)** Representative histograms (left) and quantification (right) of relative cell size, based on forward scatter area (FSC-A) assessment, and cellular granularity, based on side scatter area (SSC-A) assessment (n=15; mean ± SEM; **p<0.01 by paired Student’s t-test).

### atRA attenuates IFN-γ production of NK cells

To explore the effect of atRA on NK cell effector function, we assessed their cytokine and chemokine expression. atRA-treated NK cells displayed reduced transcript amounts of Ifng, while Csf2, Ccl5, and Ccl1 expression was elevated compared to controls (Fig. S4A). Therefore, we first analyzed the ability of atRA-treated NK cells to produce IFN-γ in response to various stimuli. When stimulated with IL-12 and IL-18, which synergistically induce IFN-γ release by NK cells [9], the cytokine amount and percentage of IFN-γ-producing atRA-treated NK cells was lower compared to controls (Fig. 3A). When activated by cross-linking of the activating receptors NK1.1, NKG2D, NKp46, or DNAM-1, atRA-treated NK cells also produced less IFN-γ compared to controls (Fig. S4B). Similar results were obtained when NK cells were incubated with tumor cells, RMA-S and YAC-1 lymphoma (Fig. S4C). These results imply that the exposure to atRA downregulates IFN-γ production by NK cells in response to various stimuli.

**Figure 3.**
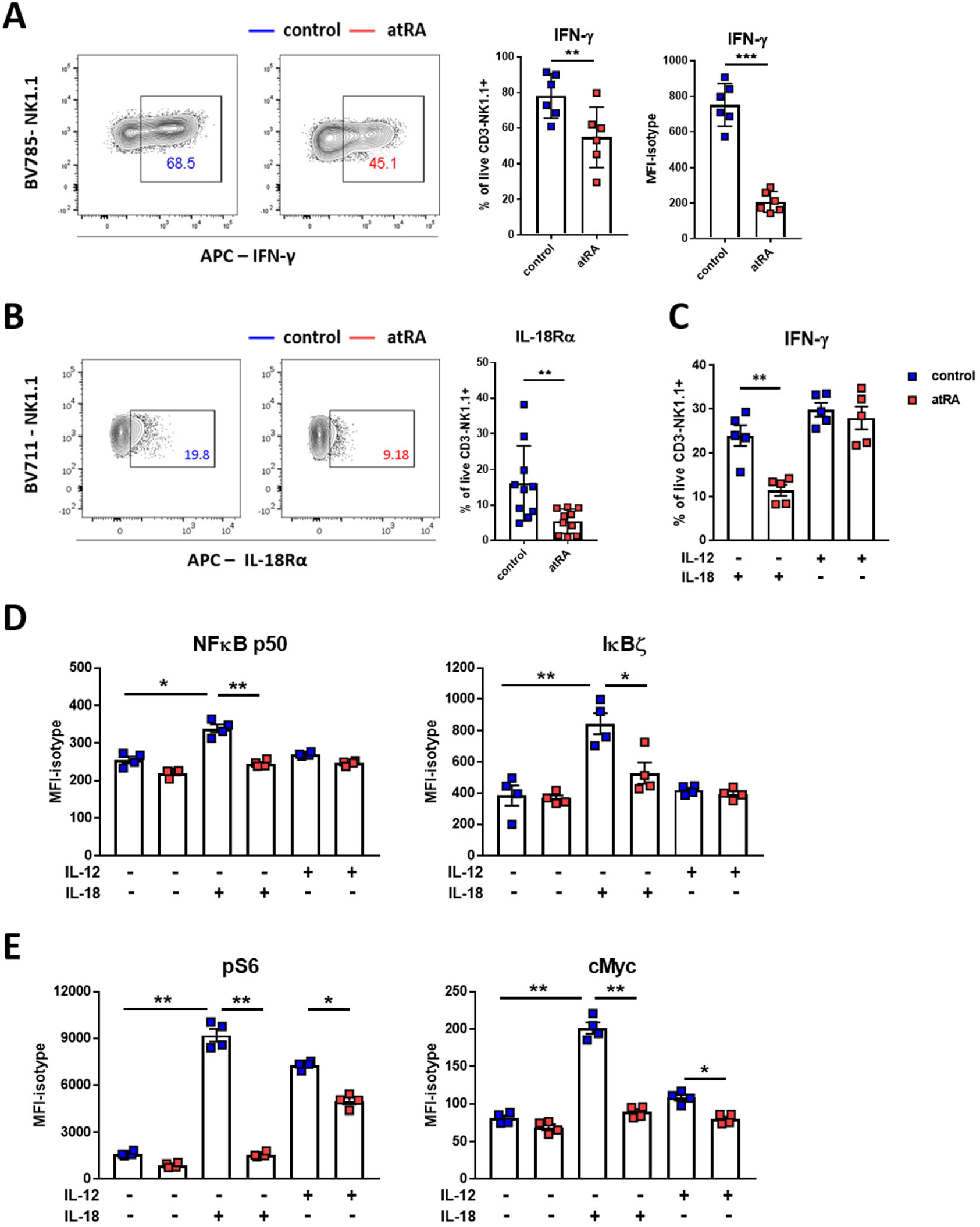
atRA-treated NK cells produce less IFN-γ and display lower activation of mTORC1 and NFκB pathway upon stimulation with IL-18. **(A)** NK cells were cultured with 1 µM of atRA (or DMSO as a control solvent) in the presence of IL-2 for 7 days and stimulated with IL-18 and IL-12 for 4 hours. Representative contour-plots (right) and quantification (left) of IFN-γ-expressing NK cells, and the produced IFN-γ amount in response to IL-12 and IL-18 (n=6; mean ± SEM; **p<0.01, ***p<0.001 by paired Student’s t-test) **(B)** NK cells were cultured with 1 µM of atRA (or DMSO as a control solvent) in the presence of IL-2 for 7 days. Representative contour-plots (left) and frequencies of IL-18Rα-expressing NK cells (right, n=10; mean ± SEM; *p<0.05 by paired Student’s t-test). **(C)** Upon culture with atRA or DMSO, cells were stimulated with IL-18 or IL-12 for 4 hours and analyzed by flow cytometry. The percentage of IFN-γ-expressing NK cells in response to IL-18 or IL-12 (n=5; mean ± SEM; **p<0.01 by paired Student’s t-test). **(D-E)** NFκB p50 and IκBζ (D), and phosphorylated (p)-S6 and cMyc (E) expression by stimulated NK cells (n=4; mean ± SEM; *p<0.05, **p<0.01 by Tukey One Way-ANOVA).

atRA-treated NK cells comprised less IL-18Rα-expressing cells compared to controls (Fig. 3B), which correlated with reduced IFN-γ production in response to IL-18, but not to IL-12 (Fig. 3C). To further investigate the mechanism of reduced IL-18-induced IFN-γ production, we analyzed the phosphorylation of proteins involved in the NFκB pathway, which mediates the responses downstream of IL-18R [27], and was transcriptionally affected in NK cells treated by atRA (Fig 1C). IL-12 and IL-18 were reported to support the production of IFN-γ via the STAT4 signaling pathway [28], and NFκB target gene - IκBζ [29], respectively. NK cells increased the expression of NFκB p50 and IκBζ upon stimulation with IL-18, but not with IL-12 (Fig. 3D and Fig. S5A), and the treatment with atRA reduced the IL-18-induced IκBζ and NFκB p50 expression. On the contrary, the phosphorylation of STAT4 was not altered by pre-exposure of NK cells to atRA (Fig. S5B). Similarly, the phosphorylation of NFκB p65 and kinase IKK α/β upon stimulation with IL-12 or IL-18 was unchanged (Fig. S5C). Furthermore, we investigated the activation of mTORC1, which drives glycolysis and lipid synthesis by regulating the surface expression of nutrient transporters [30], using the phosphorylation of ribosomal protein S6 as a read-out, and the expression of cMyc, which is required for cytokine-induced metabolism and effector functions in NK cells [31]. Phosphorylation of S6 was induced with either IL-12 or IL-18, while cMyc expression increased in NK cells only upon exposure to IL-18 (Fig. 3E and Fig. S5A). However, atRA-exposed NK cells failed to activate mTORC1 and to increase cMyc expression in response to IL-18 (Fig. 3E and and Fig. S5A). Altogether, atRA-treated NK cells produced less IFN-γ compared to control NK cells in response to IL-18, but not IL-12, and this correlated with reduced activation of NFκB-IκBζ and the mTORC1 pathway, and the reduced expression of cMyc.

### atRA-treated NK cells favor immature phenotype of DCs

It was previously reported that NK cells release IFN-γ while interacting with DCs, and promote DC maturation and the expression of antigen presenting and co-stimulatory molecules on DCs. Reciprocally, DCs produce IL-12 and IL-18 during the interaction with NK cells [9, 32]. To address if atRA-treated NK cells regulate DC phenotype, we co-cultured atRA-treated or control NK cells with bone marrow-derived dendritic cells (BM-DCs), and analyzed their maturation and survival. In accordance with literature, DCs increased expression of CD86 and MHC II upon co-culture with control NK cells, while atRA-treated NK cells showed a reduced ability to upregulate the expression of these molecules on DCs (Fig. 4A). Accordingly, we detected lower amounts of IFN-γ (Fig. 4B) and lower percentages of apoptotic DCs (Fig. 4C) in the co-culture with atRA-treated NK cells, indicating that NK cells exposed to atRA favor immature DC phenotype, which correlates with their lower ability to secrete IFN-γ in response to DCs.

**Figure 4.**
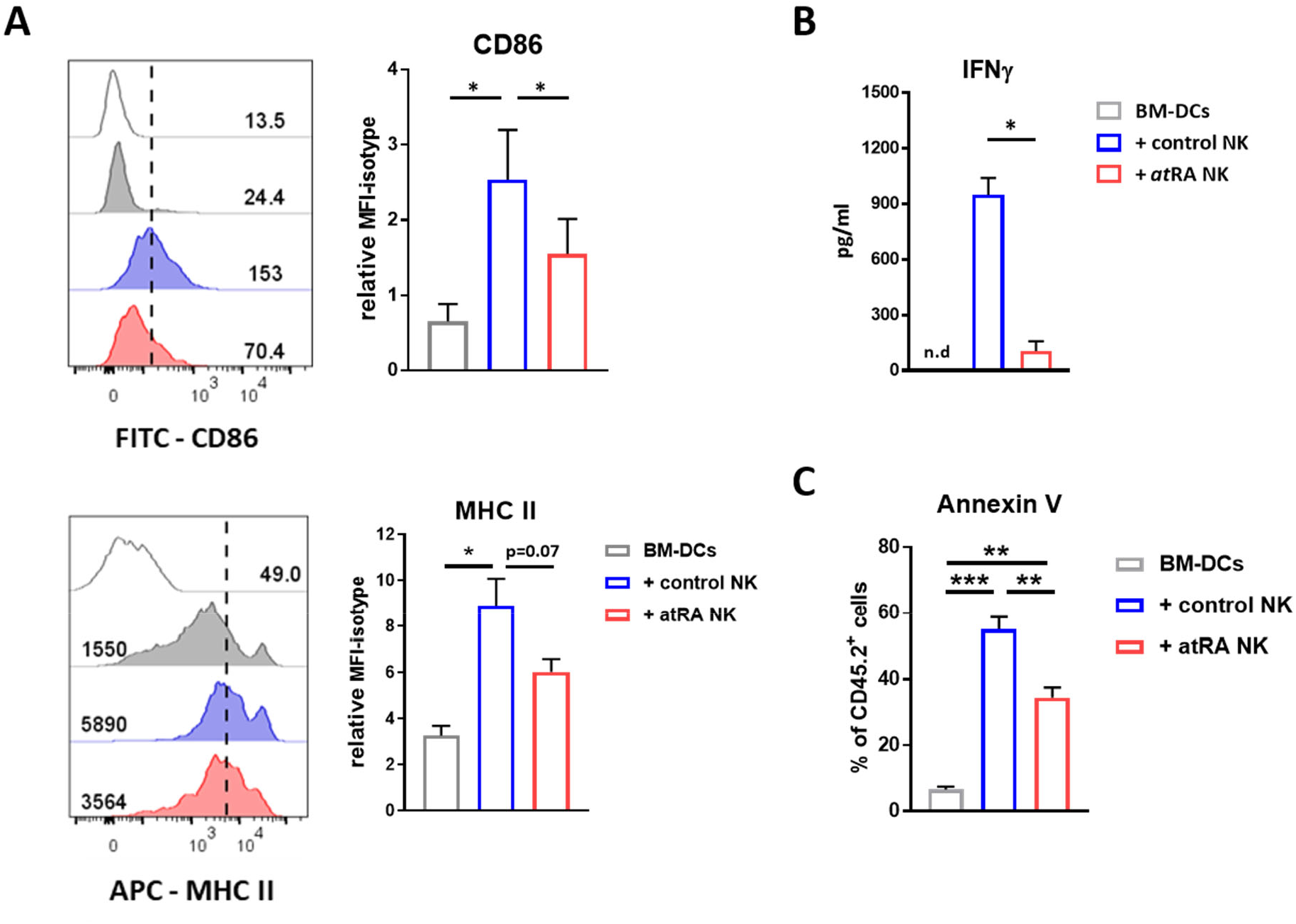
atRA-treated NK cells favor immature BM-DC phenotype. **(A)** Representative histograms (left) and quantification (right) of CD86 and MHC II expression by BM-DCs cultured with atRA-treated or control NK cells for 24 hours (n=4; mean + SEM; *p<0.05 by Tukey One Way-ANOVA). **(B)** Quantification of secreted IFN-γ in the DC-NK cell co-culture (n=3; mean + SEM; *p<0.05 by paired Student’s t-test). **(C)** The percentage of apoptotic (Annexin V-labeled) BM-DCs after 6 hours of co-culture with NK cells (n=2-4; mean + SEM. *p<0.05, **p<0.01 by Tukey One Way-ANOVA).

### atRA-treated NK cells support regulatory T cell differentiation

NK cells and NK cell-derived IFN-γ were also reported to support the differentiation of Th1 cells [11, 33]. To determine whether atRA affects the ability of NK cells to regulate T cell differentiation, we co-cultured atRA-treated or control NK cells with naïve CD4^+^ T cells in Th1-, Th17-, or Treg-polarizing conditions. In Th1-polarizing conditions, the expression of the transcription factor T-bet and IFN-γ by CD4^+^ T cells was not affected by the NK cells, irrespective of their pre-treatment (Fig. S6A). However, control NK cells reduced the ability of naive T cells to differentiate into FoxP3-expressing Tregs, while the frequency of differentiated Tregs was not affected by atRA-treated NK cells (Fig. 5A). In Th17-polarizing conditions, the majority of CD4^+^ T cells acquired the expression of the transcription factor Rorγt, which was not affected by NK cells (Fig. 5B). Among Rorγt^+^ CD4^+^ T cells, a subpopulation of cells co-expressed FoxP3, and the percentage of FoxP3^+^ CD4^+^ Th17 cells increased in the presence of atRA-treated NK cells, compared to both control NK cells and to T cells cultured alone (Fig. 5B).

**Figure 5.**
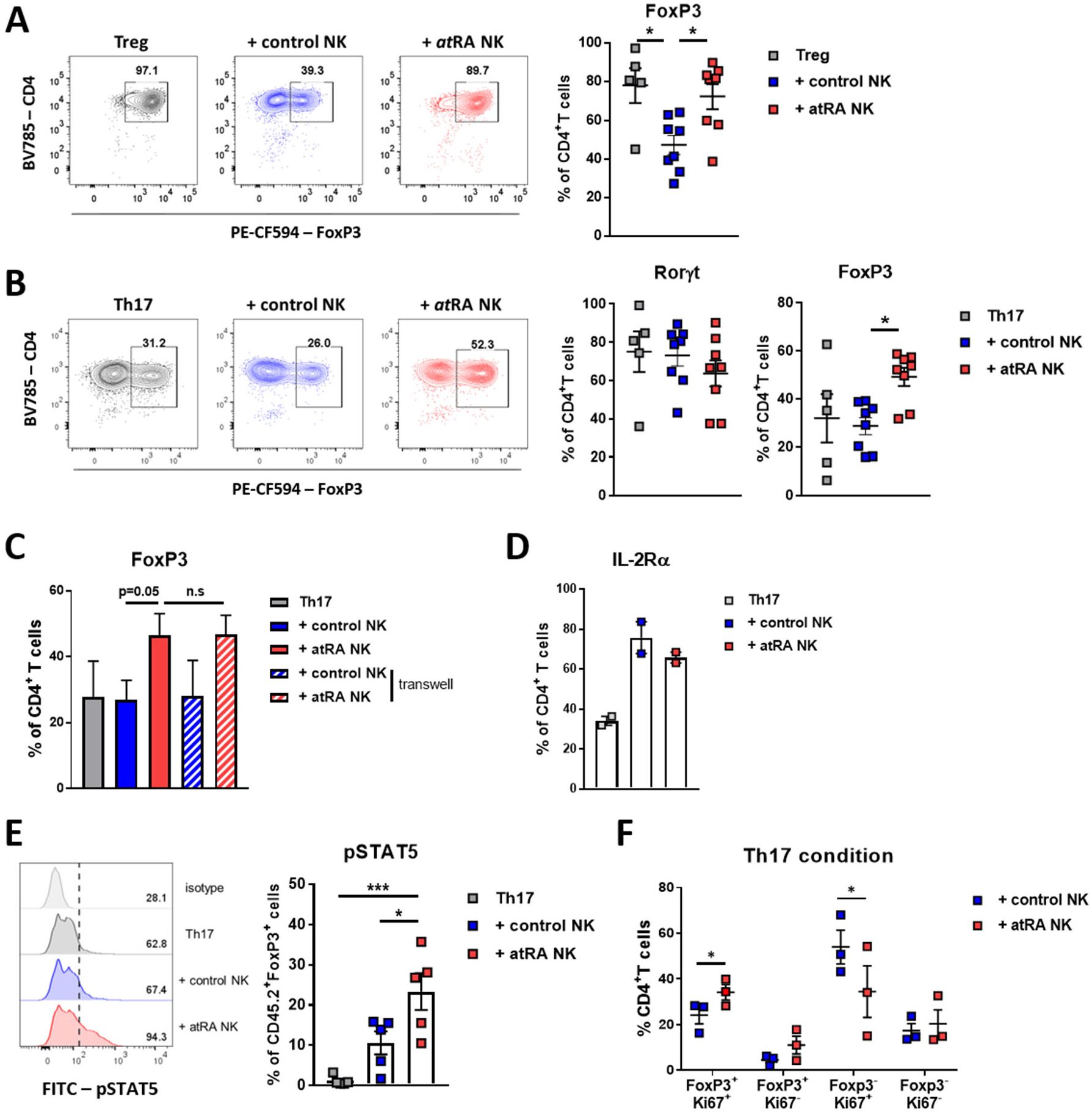
atRA-treated NK cells promote the differentiation and proliferation of FoxP3-expressing T cells. **(A-B)** CD45.1^+^ NK cells were treated with 1 µM of atRA (or DMSO as a solvent control) in the presence of IL-2 for 5 days, and then co-cultured with freshly-isolated naïve CD45.2^+^ CD4^+^ T cells in Treg or Th17-polarizing conditions. T cells were analyzed for the expression of Rorγt, FoxP3, Ki67, CD25, and phosphorylated (p)-STAT5 by flow cytometry. Representative contour-plots (left) and frequencies (right) of FoxP3- and/or Rorγt-expressing CD4^+^ T cells in (A) Treg-polarizing conditions (n=5-8) and (B) Th17-polarizing conditions (n=5-8). The graphs indicate mean ± SEM; *p<0.05 by Tukey OneWay-ANOVA. **(C)** Frequencies of FoxP3-expressing CD4^+^ T cells cultured in Th17-polarizing conditions with NK cells separated using permeable transwell membrane (n=3, mean + SEM; n.s. not significant and p=0.05 by Tukey OneWay-ANOVA). **(D)** Percentage of IL-2Rα-expressing CD4^+^ T cells upon co-culture with NK cells (n=2). **(E)** Representative histograms (left) and quantification (right) of p-STAT5 in CD45.2^+^ T cells cultured alone, with control NK cells, or with atRA-treated NK cells (n=5; mean ± SEM. *p<0.05, ***p<0.0001 by Tukey OneWay-ANOVA). **(F)** Frequency of FoxP3^+^ Ki67^+^ and FoxP3^neg^ Ki67^+^ CD4^+^ T cells upon co-culture with NK cells (n=3; mean ± SEM. *p<0.05 by paired Student’s t-test).

To address the mechanism by which atRA-treated NK cells supported the differentiation of FoxP3^+^ Th17 cells, we first prevented a direct NK cell-to-T cell contact using transwell-chamber system. In this setup, FoxP3 expression by CD4^+^ T cells was not affected (Fig. 5C), indicating that soluble molecules supported the differentiation of FoxP3^+^ CD4^+^ cells. Among these, IL-2-receptor signaling via STAT5 was shown to be required for the Treg development [34, 35], and adenosine, IL-10, CCL1, and CCL5 were reported to support Treg differentiation [36–39]. Since we detected increased expression of adenosine-metabolizing enzymes (CD73, CD39, and CD38; Fig. 1F), and the Ccl1 and Ccl5 transcripts (Fig. S4A) in NK cells exposed to atRA, we analyzed whether these molecules regulate regulatory cell differentiation in our NK-Th17 co-culture system. Neither antagonist-mediated blockade of the adenosine receptor A2AR, nor the mAb-mediated neutralization of IL-10, CCL1, or CCL5, affected FoxP3 expression by CD4^+^ T cells upon co-culture with NK cells (Fig. S6B-D). However, although the percentage of IL-2Rα-expressing T cells was comparable (Fig. 5D), the frequency of phosphorylated-STAT5^+^-expressing T cells was higher in co-cultures with atRA-treated compared to controls (Fig. 5E). This correlated with a higher percentage of Ki67^+^ FoxP3-expressing CD4^+^ T cells (Fig. 5F and Fig. S6E), suggesting that NK cells exposed to atRA could support the proliferation of FoxP3^+^ CD4^+^ T cells via the phosphorylation of STAT5, downstream of IL-2 receptor engagement.

## Discussion

Vitamin A is an essential lipid-soluble nutrient, which promotes tolerogenic immunity in the gastrointestinal tract [40]. Primary role of vitamin A in gut is to create microenvironment in which immune cell functions are skewed towards promoting immune regulation. For, example, RA-imprinted DCs were shown to support the Treg differentiation, and IL-10 production [41, 42], and the expression of gut-homing receptors on B cells and Tregs [41, 43]. Similarly, ILC3s were reported to increase the expression of gut-homing receptors upon the exposure to RA [25, 44, 45]. In mice fed with vitamin A-deficient diet, the ratio of IL-22-producing ILC3s to ILC2s declined, resulting in an increased host susceptibility to bacterial infection [46]. Our data show that atRA does not increase CCR9 and CD62L expression in NK cells, indicating that vitamin A might not imprint a “classical” lymph node- and/or gut-homing phenotype in NK cells, and might rather regulate NK cell phenotype and function in vitamin A-enriched microenvironments other than gut. The organ with highest amounts of stored vitamin A and its derivatives is the liver [47]. In the steady-state, the liver tissue is characterized by tolerogenic immunoregulatory environment. T cells primed in the liver predominantly differentiate towards a regulatory phenotype, in a process that is orchestrated by local antigen presentation [48], inhibitory checkpoints molecules [49] and microenvironmental cytokines [50]. In the gut mucosa, Treg and Th17 cells are enriched to restrain inflammatory responses and protect the host from infection, respectively. Despite their different functions, they share similar differentiation modules, tightly regulated by microenvironmental factors, such as IL-6, TGF-β, IL-1β, IL-21, short-chain fatty acids and RA [51–53]. It was reported that the outcome of naive T cell differentiation towards Treg or Th17 effector cell depends on relative concentrations of key cytokines regulating these cell fates [54, 55]. RA supports Treg differentiation by enhancing TGF-β over IL-6 signaling [56]. Th17 cells appear to be particularly “plastic”, showing high adaptability to local cytokine milieus. They can acquire both IFN-γ production and Th1-like phenotype, or to co-express FoxP3 and IL-10, favoring immune regulation [57, 58]. However, it is not clear how FoxP3^+^ cells develop within the Th17 differentiation program. Gagliani et al. also reported the existence of intestinal “ex-Th17 cells” under homeostasis, that irreversibly lost the ability to produce IL-17, and acquired Tr1-like phenotype [59]. Although key cytokine players that shape the Th17-Treg plasticity and trans-differentiation have been identified, the contribution of other environmental factors, including interactions with neighboring cells, especially in tissues other than gut, is still elusive. We show that in both Th17- and Treg-polarizing conditions, atRA-treated NK cells supported the differentiation and/or proliferation of FoxP3-expresing CD4^+^ T cells. This effect was mediated by soluble factors, and correlated with increased phosphorylation of STAT5 in T cells. STAT5 is activated upon cytokine-receptor engagement, including receptors for IL-3, IL-4, IL-5, GM-CSF and cytokines signaling via the common γc chain [60, 61]. As STAT5 signaling and IL-2 were reported to be essential for the development of Tregs [34, 35], and suppression of Th17 program [62], IL-2-IL-2R might drive atRA-mediated effects in co-culture of NK cells with T cells. In addition, our findings also indicate the potential of atRA-treated NK cells to produce adenosine, due to co-expression of the adenosine-generating enzymes CD73, CD39 and CD38. Adenosine is immunosuppressive mediator derived from extracellular ATP that accumulates in pathological conditions, such as in hypoxic solid tumor tissue [63]. Stimulation of adenosine receptor A2AR in naive CD4^+^ T cells was shown to promote Treg development and FoxP3 expression [64]. Although A2AR blockade did not indicate its role in our co-culture system, this axis might synergistically support immunoregulatory NK cells functions of NK cells in vitamin A-enriched tissues. In the liver, where NK cells comprise the majority of lymphocytes [65], the adenosine pathway plays an important roles in regulating inflammation and immune responses in liver disease [66, 67]. Liver tissue injury leads to concomitant release of intracellular ATP from damaged and necrotic cells and to the depletion of vitamin A storage from hepatic stellate cells (HSCs) [17]. Released retinol can be converted to its active form(s) by HSCs and by neighboring stromal and parenchymal cells [68], thus becoming available for ongoing tissue responses, while ATP is being converted to adenosine. How these microenvironmental milieus regulate NK cell responses has still not been addressed.

Our results show that exposure to atRA primarily reduces NK cell pro-inflammatory responses, such as IFN-γ production. IFN-γ is an important effector cytokine in both innate and adaptive immunity, which can be produced by both NK cells and activated T cells [69, 70]. atRA was shown to downregulate the production of IFN-γ by CD8^+^ T cells (Van et al., 2009), as well as by NKT cells in mice with concanavalin A - induced hepatitis [71]. An important role of IFN-γ during early immune response activation is to promote differentiation and maturation of monocytes and DCs [72, 73], and NK cell-derived IFN-γ was shown to support the expression of co-stimulatory molecules and antigen-presenting machinery in maturing DCs [9, 10]. At the same time, by eliminating immature DCs, NK cells can assure the generation of an activated mature DC pool to drive efficient T cell priming. We show that atRA-treated NK cells secreted less IFN-γ during co-culture with DCs, which correlated with lower expression of co-stimulatory molecules, and higher survival of immature DCs. Several receptor-ligand interaction, as well as soluble factors were reported to contribute to NK-DC cross-talk resulting in maturation and/or cytotoxicity [7]. atRA-mediated reduction of IFN-γ production by NK cells seems to comprise a rather general mechanism that operates downstream of various stimuli, including cytokines, activating receptor triggering or encounter of tumor target cells. The underlying mechanism targets two signaling axes that regulate NK cell effector functions and cellular fitness. First, our results show that NK cells exposed to atRA were unable to efficiently engage mTORC1 and increase cMyc expression in response to IL-18. mTORC1 and cMyc are metabolic regulators that increase glycolysis rate and support pro-inflammatory effector responses in T cells [74–76]. NK cells require mTORC1 for development, as well as for the IL-15-induced IFN-γ production by elevating glycolytic rate [30]. cMyc protein expression in NK cells was shown to be regulated by glutamine availability and SLC7A5-dependent amino acid transport [31]. cMyc-deficient NK cells expanded in presence of IL-15, failed to undergo blastogenesis, to increase mitochondrial mass and metabolic flux, and produced less IFN-γ in response to IL-2 and IL-12 [31]. Similarly, atRA-exposed NK cells displayed reduced rates of both glycolysis and oxidative phosphorylation compared to controls, reduced cell size, and reduced mitochondrial mass and membrane potential. Although previous studies reported the impact of atRA on mitochondria and cellular metabolism that appears to be cell- and organ-specific [77–79], it remains to be investigated if atRA mediates these effects through regulation of cMyc.

In addition to mTORC1 and cMyc, the attenuated IFN-γ production by atRA-treated NK cells correlated with lower expression of genes involved in the NFκB pathway, as well as lower expression of NFκB p50 in response to stimulation of the IL-18 receptor. Stimulation with IL-12 was not affected by atRA, indicating that atRA modulated preferentially NFκB-dependent responses. The NFκB signaling cascade operates downstream of several other receptors expressed by NK cells, including NKG2D, DNAM-1, CD16 or Ly49H [80], which might explain the reduced IFN-γ production when atRA-exposed NK cells were triggered via these receptors, or with tumor cells previously reported to express their cognate ligands [81–83]. NFκB was reported to directly bind to the *Ifng* promoter [84]. However, NFκB also induces the expression of transcription factor Iκbζ that also regulates *Ifng* transcription [29]. We have previously reported that in hypoxic conditions, IFN-γ production by NK cells was attenuated through HIF-1α-mediated inhibition of IL-18-NFκB-Iκbζ pathway [16]. In addition, Iκbζ-deficient mice display reduced IFN-γ production by NK cells, but not other cells [29], indicating that along with metabolic reconfiguration, the NFκB-Iκbζ axis might be the predominant checkpoint for regulating IFN-γ release by NK cells.

In summary, we identified a novel role of vitamin A metabolite, atRA, in regulating NK cell phenotype, metabolism and function. atRA-exposed NK cells display a reduced size, mitochondrial mass and membrane potential, associated with reduced metabolic rates. They favor an immunoregulatory microenvironment through a lower ability to remove immature DCs and to produce IFN-γ in response to wide range of stimuli. In addition, atRA-treated NK cells support FoxP3^+^ T cell differentiation and/or proliferation. Therefore, in vitamin A-enriched tissue environments, NK cells can contribute to the maintenance of homeostasis, or to regulation and/or resolution of inflammatory responses in pathological settings.

## Materials and methods

### Mice

Rag2-deficient mice were bred and maintained at the animal facility of Medical Faculty Mannheim, University of Heidelberg. C57BL6/N mice were purchased from Janvier. Both male and female mice were used for experiments (8-40 weeks of age). All animal experiments were approved by the local authorities.

### Preparation of single-cell suspension from spleen and bone marrow

Single-cell suspensions were obtained from spleens after red blood cell lysis with buffered chloride potassium phosphate solution (ACK buffer) for 2-5 min at RT. Single-cell suspensions from bone marrow were obtained by flushing femurs and tibias with PBS. Subsequently, cells were collected by centrifugation, and red blood cells were lysed with ACK buffer.

### NK cell isolation and culture

NK cells were enriched from splenic single-cell suspensions of Rag2-deficient mice after removal of adherent cell fraction by short culture at 37°C. Cells were then cultured for 5-7 days in RPMI-1640 medium (Sigma-Aldrich) supplemented with 10% heat-inactivated FCS, 2 mM glutamine, 1% non-essential amino acids, 1 mM sodium pyruvate, 100 U/ml Penicillin/Streptomycin, 50 µM β-mercaptoethanol (all from GIBCO) and 1700 U/ml recombinant human IL-2 (NIH). All-*trans* retinoic acid (atRA; Sigma-Aldrich) was dissolved in dimethyl sulfoxide (DMSO; Sigma-Aldrich) and added at the concentrations indicated in the Results. Equivalent volumes of DMSO were used as control.

### Tumor cell lines

The mouse lymphoma cell lines RMA-S and YAC-1 were cultured in RPMI-1640 medium supplemented with 10% FCS and 100 U/ml Penicillin/Ampicillin. The mouse melanoma cell line B16 was cultured in Dulbecco’s Modified Eagle Medium (DMEM; Sigma-Aldrich) supplemented with 10% FCS and 100 U/ml Penicillin/ Streptomycin.

### *In vitro* stimulation of NK cells

After 7 days of culture with atRA, NK cells were cultured for 5 hours with RMA-S, YAC-1 or B16 cells (1:1), or stimulated with cytokines (1 ng/mL of IL-12 and 10 ng/mL of IL-18), plate-bound antibodies (Abs) anti-NK1.1 (clone PK136; Biolegend), anti-NKG2D (clone A10; Biolegend), or anti-NKp46 (polyclonal goat IgG; R&D Systems), or plate-bound CD155-Fc chimera protein (CD155-IEGRMDP-Mouse IgG2a; R&D Systems). For plate coating with Abs or Fc, flat-bottomed 96-well plates were incubated overnight at 4°C with 10 µg/ml Ab or 2 µg/ml chimera protein solution in PBS. Stimulation was performed in the presence of fluorophore-labeled anti-CD107a mAb (clone 1D4B; Biolegend) or the isotype-matched mAb control. GolgiStop™ (BD Biosciences) was added 4 hours before the end of stimulation.

### Flow-cytometry and cell sorting

Cells were incubated with Fc-receptor-blocking reagent (10% supernatant of αCD16/CD32-producing hybridoma 2.4G2), followed by incubation with fluorochrome-conjugated monoclonal antibodies directed against CD45.1 (clone A20), CD45.2 (clone 104), CD4 (clone GK1.5), CD3ε (clone 145-2C11), NK1.1 (clone PK136), NKG2D (clone CX5), NKp46 (clone 29A1.4, CD226 (clone 10E5), CD200R (clone OX-2R), CD11b (clone M1/70), CD27 (clone LG.3A10), LAG3 (clone C9B7W), TRAIL (clone N2B2), CD62L (clone MEL-14), KLRG1 (clone 2F1), CD39 (clone Duha59), CD73 (clone TY/11.8), CD38 (clone 90), IL-18Rα (clone A17071D), CD86 (clone GL-1), CD25 (clone PC61) (all from Biolegend), and I-A/I-E (clone M5/114.15.2; BD Biosciences). For the detection of cytokines and transcription factors, antibodies specific for IFN-γ (clone XMG1.2; Biolegend), Ki67 (clone 16A8; Biolegend), IκBζ (clone LK2NAP; Invitrogen), cMyc (clone D84C12; Cell Signaling Technology), NFκB-p50 (clone D4P4D; Cell Signaling Technology), Eomes (clone X4-83; BD Bioscience), FoxP3 (clone FJK-16s; eBiosciences), Rorγt (clone Q31-378; BD Biosciences) and T-bet (clone 4B10; Biolegend), and eBioscience™ Foxp3/Transcription Factor Staining Buffer Set (Thermo Fisher) were used according to the manufacturer’s instructions. For the detection of phosphorylated proteins, STAT4 (clone D2E4), NFκB-p65 (clone 93H1), S6 (clone D57.2.2E), IKKα/β (clone 16A6), and STAT5 (polyclonal rabbit IgG) (all from Cell Signaling Technology), BD Cytofix/Cytoperm™ and BD Phosflow™ buffer (BD Biosciences) were used according to the manufacturer’s instructions. 7-aminoactinomycin D (7-AAD; BD Bioscience) or Zombie Aqua™ (Biolegend) were used to exclude dead cells. For the analysis of lipid uptake and mitochondria, Bodipy™ FL C16 (Invitrogen) or CellROX™ Deep Red dye, MitoTracker™ Green FM dye, and MitoProbe™ tetramethylrhodamine methyl ester (TMRM) dye (all from Thermo Fischer) were used, according to the manufacturer’s instructions. For the detection of apoptosis, Annexin V Apoptosis Detection Kit (Biolegend) was used. Flow-cytometry was performed with LSRFortessa™, and data were analyzed using FlowJo™ software (both from BD Biosciences). For the gene expression analysis, NK cells (NK1.1^+^NKp46^+^TRAIL^neg^CD200R^neg^) were sorted from pre-gated live/single/CD45^+^ splenocytes of Rag2-deficient mice using BD FACSAria™ Fusion (BD Biosciences).

### Measurement of oxygen consumption rate (OCR) and extracellular acidification rate (ECAR)

The analysis was performed using Seahorse™ XF HS Mini Analyzer (Agilent). After 7 days of culture, NK cells were resuspended in XF RPMI medium (Agilent) containing 10 mM glucose, 2 mM L-glutamine, and 1 mM sodium pyruvate. The OCR and ECAR were analyzed in response to 1.5 μM oligomycin, 2.5 μM BAM15 ((2-fluorophenyl){6-[(2-fluorophenyl)amino](1,2,5-oxadiazolo[3,4– e]pyrazin-5-yl)}amine), and 0.5 μM Rotenone and Antimycin A (all from Agilent), according to the manufacturer’s recommendations.

### Co-culture of NK cells and DCs

To obtain DCs, bone-marrow cells of C57BL6/N mice were cultured in DMEM medium supplemented with 10% FCS, 2 mM glutamine, 1% non-essential amino acids, 1 mM sodium pyruvate, 100 U/ml Penicillin/Streptomycin (all from GIBCO) and GM-CSF (10% supernatant of GM-CSF-producing X6310 cell line). NK cells were co-cultured with DCs at ratio 1:2 for the time indicated in the Result sections. Collected supernatants were assessed for cytokine content using ELISA MAX™ Standard Set Mouse IFN-γ (Biolegend). DCs were analyzed for the expression of CD86 and MHC class II using flow cytometry. DC apoptosis was assessed by measuring Annexin V binding to the cell surface.

### Differentiation of CD4^+^ T cells

Naïve CD4^+^ T cells were isolated from spleens of C57BL6/N mice using magnetic cell sort (Miltenyi). The purity of naïve T cells (CD45^+^CD3ε^+^CD4^+^CD62L^+^CD44^neg^) was ≥ 90%. T cells were activated with plate-bound anti-CD3ε (clone 145-2C11; Biolegend) and 0.5 µg/ml of soluble anti-CD28 (clone 37.51; Biolegend) mAbs. Following combinations of cytokines and antibodies were used for the differentiation of T cells: for Th1 cells - 1 µg/ml anti-IL-4 (clone 11B11; Biolegend), 30 U/ml rhIL-2 (NIH), and 10 ng/ml IL-12 (Peprotech); for Th17 - 1 µg/ml anti-IFN-γ (clone R4-6A2), 1 µg/ml anti-IL-2 (clone JES6-1A12), 1 µg/ml anti-IL-4 (clone 11B11) (all from Biolegend), 20 ng/ml IL-6, 20 ng/mL IL-1β (both from Peprotech), 1 ng/ml TGFß1, and 10 ng/ml IL-23 (both from Biolegend); for Treg - 1 µg/ml anti-IFN-γ (clone R4-6A2), 1 µg/ml anti-IL-4 (clone 11B11) (both from Biolegend), 30 U/ml rhIL-2 (NIH), and 2 ng/ml TGF-ß1 (Biolegend). NK cells treated with atRA for 5 days were collected and cultured with naïve T cells at a ratio of 1:1 for 2 days. To measure T cell cytokine production, cells were stimulated with 50 ng/mL of Phorbol myristate acetate (PMA) and 1 µM of ionomycin. In some experiments, 5 μM of ZM 241385, 5 μM of SCH 58261 (both A2AR agonists, Tocris), 7.5 μg/mL of anti-IL-10 antibody (clone JES5-2A5, BioXCell), 4 μg/mL of anti-CCL5 Ab (polyclonal goat IgG; R&D Systems), or 4 μg/mL of anti-CCL1 Ab (clone 148113; R&D Systems), was added to the cultures.

### RNA isolation and quantitative PCR

After 7 days of culture with atRA or equivalent volume of DMSO, NK cells were lysed with RLT buffer containing 1% of β-mercaptoethanol, and RNA extraction was performed using RNeasy^®^ Mini Kit (Qiagen). Genomic DNA was removed by TURBO DNA-free™ kit (ThermoFisher). RNA concentration was determined using plate reader Infinite 200 pro (TECAN). cDNA was synthesized from a total of 400 ng of RNA, using RT^2^ First Strand Kit (Qiagen). For the analysis of relative mRNA expression, RT^2^ Profiler™ PCR array mouse cytokines & chemokines (Qiagen) was used according to manufacturer’s instructions. mRNA expression was calculated using 2^-ΔCP^ method, relative to b2m transcript.

### Gene expression analysis

NK cells isolated from spleens of Rag2^-/-^ mice using flow-cytometric sort were cultured with atRA or equivalent volume of DMSO for 7 days. Gene expression profiling was performed using Clariom™ D Mouse Arrays (Thermo Fisher). Biotinylated antisense cDNA was prepared according to the standard labeling protocol with the GeneChip^®^ WT Plus Reagent Kit and the GeneChip^®^ Hybridization, Wash and Stain Kit (both from Thermo Fisher). Chip hybridization was performed using GeneChip Hybridization oven 640, then dyed in the GeneChip Fluidics Station 450, and scanned with a GeneChip Scanner 3000 (all from Affymetrix). Raw fluorescence intensity values were normalized by applying quantile normalization and RMA background correction. A custom CDF version 22 with ENTREZ based gene definitions was used to annotate the arrays [85]. Differentially expressed transcripts were obtained by One Way-analysis of variance (ANOVA) using a commercial software package SAS JMP15 Genomics, version 10, from SAS (SAS Institute). Parameters applied were a false positive rate of α = 0.05 with FDR correction as the level of significance. Data are visualized using EnhancedVolcano (https://github.com/kevinblighe/EnhancedVolcano) and Pheatmap package in R (https://github.com/raivokolde/pheatmap). For pathway enrichment analysis, following criteria were used to choose specifically expressed genes: fold change > 1.5 or < −1.5 and adjusted p-value < 0.05. Pathway enrichment analysis was based on the pathway collection in Kyoto Encyclopedia of Genes and Genomes (KEGG) (https://www.genome.jp/kegg/pathway.html).

### Quantification and statistical analysis

Experimental results are shown as mean ± SEM; n represents numbers of experiments, and is specified in Figure Legends. The statistical significance was assessed with the paired Student’s t-test for normally distributed data, Welch’s t-test for data with unequal variances, One Way-ANOVA followed by Tukey’s post-hoc test for multiple testing, all using with GraphPad Prism software (GraphPad Software). The differences between experimental groups were considered significant when *p < 0.05, **p < 0.01, ***p < 0.001, ****p <0.0001.

## Supporting information

Jeong_etal_SupplementalData

## Abbreviation

A2AR: adenosine A2A receptor
atRA: all-*trans* retinoic acid
BM-DCs: bone marrow-derived dendritic cells
CD: cluster of differentiation
CCL: chemokine (C-C motif) ligand
DCs: dendritic cells
DMSO: dimethyl sulfoxide
ECAR: extracellular acidification rate
FCS: fetal calf serum
GM-CSF: granulocyte-macrophage colony-stimulating factor
ILC3s: group 3 innate lymphoid cells
IκBζ: inhibitor of kappa B-zeta
IKK α/β: IκB kinase alpha/beta
IL: interleukin
IFN: interferon
mAb: monoclonal antibody
MHC: major histocompatibility complex
NK: natural killer
NIH: National Institutes of Health
NFκB: nuclear factor kappa B
OCR: oxygen consumption rate
PBS: phosphate buffered saline
RA: retinoic acid
ROS: reactive oxygen species
RT: room temperature
RPMI: Roswell Park Memorial Institute
STAT: signal transducer and activator of transcription
Th1: T helper type 1
Th17: T helper type 17
TNF: tumor necrosis factor
Tregs: regulatory T cells
TORC1: target of Rapamycin Complex 1

## Acknowledgments

We thank Marian Wincher, Petra Bugert, Dalyan Devran and Patrick Matei for technical support; Sophia Papaioannou for help with experiments; the animal facility of the Medical Faculty Mannheim for assistance with animal care and experiments; and Dr. Margareta Correia and Prof. Guoliang Cui for support and discussion of the data. The project was supported by the Research Training Group at Heidelberg University train4CIM (to AC); intramural funding program SEED by Medical Faculty Mannheim (to AS); grants from the German Research Foundation [SPP1937 (CE 140/2-2 to AS and AC), SFB1366 (Project number 319 394046768-SFB 1366; C02 to AC), SFB-TRR156 (B10N to AC), RTG2727 – 445549683 Innate Immune checkpoints in cancer and tissue damage (B1.2 to AC and AS), TRR179 (TP07 to AC)]; ExU 6.1.11 (to AC), German Cancer Aid grant “NK fit against AML” (73000336 to AC); a network grant of the European Commission (H2020-MSCA-MC322, ITN-765104-MATURE-NK); and by the Angioformatics platform of the European Center for Angioscience (ECAS).

## Author contributions

MJ and AS designed experiments and wrote the manuscript; MJ performed experiments and analyzed the data; JXS performed experiments; CDLT performed gene expression analysis. AS and AC supervised the study.

