## Supplementary material for "Vitamin A-treated natural killer cells reduce interferon-gamma production and support regulatory T cell differentiation": Jeong_etal_SupplementalData

### Figure S1

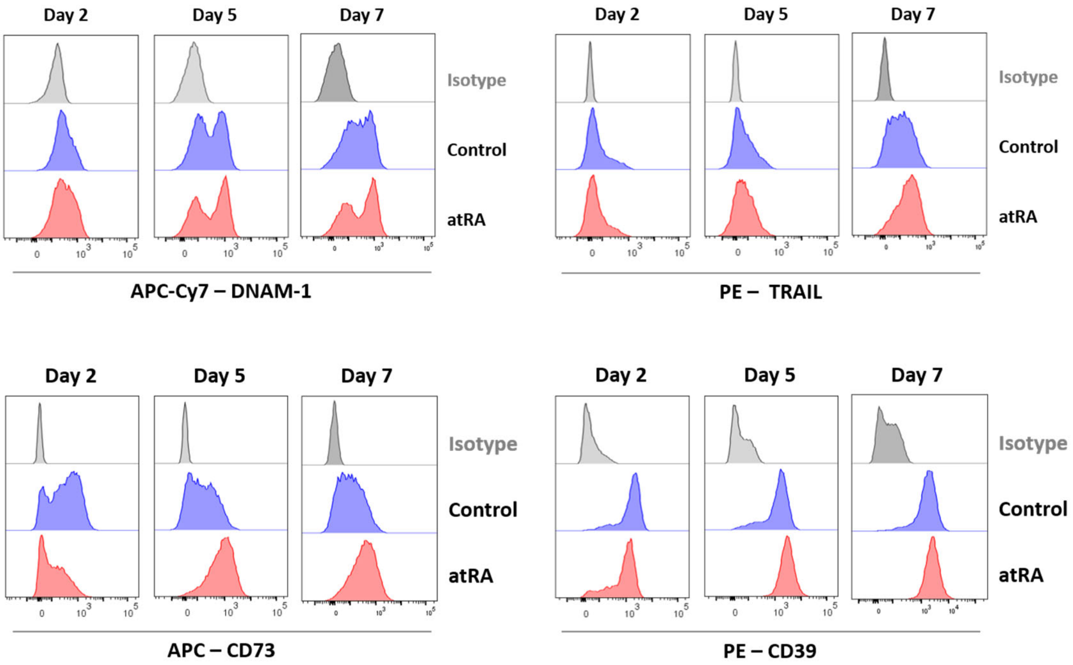


**Figure S1. atRA-treated NK cells upregulate DNAM-1, TRAIL, CD73, and CD39 in a time-dependent manner.**

NK cells were cultured with 1 µM of atRA (or the equivalent volume of DMSO as a solvent control) in the presence of IL-2 for 2, 5, or 7 days. The expression of indicated molecules was analyzed by flow cytometry. Shown are the histograms representative of four independent experiments.

**Figure S2**

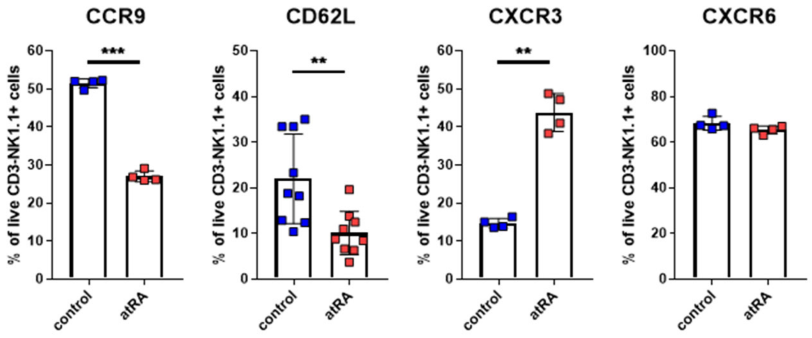


**Figure S2. Expression of CCR9, CD62L, CXCR3, and CXCR6 by atRA-treated and control NK cells.**

NK cells were cultured with 1 µM of atRA (or the equivalent volume of DMSO as a solvent control) in the presence of IL-2 for 7 days. The expression of indicated molecules was analyzed by flow cytometry (n=4-9; mean ± SEM. **p<0.01, ***p<0.001 by paired Student's t-test).

**Figure S3**

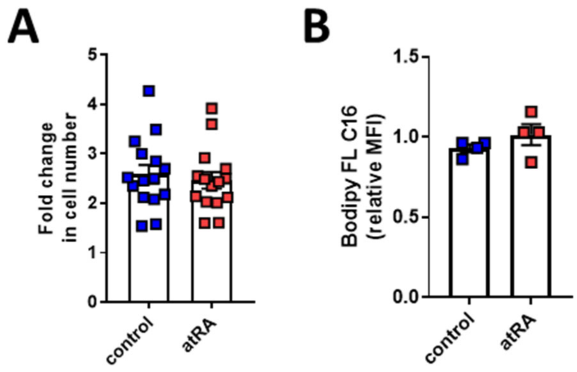


**Figure S3. atRA does not affect NK cell expansion and lipid uptake.**

**(A)** NK cells were cultured with 1 µM of atRA (or the equivalent volume of DMSO as a control solvent) in the presence of IL-2 for 7 days. Quantification of cell expansion rates at the end of culture (day 7) calculated as final cell numbers divided by cell numbers seeded at the beginning of the culture (day 0), (n=15; mean ± SEM; not significant by paired Student's t-test)

**(B)** Geometric mean fluorescence intensity (MFI) of Bodipy™ FL C16 staining analyzed by flow cytometry (n=4; mean ± SEM; not significant by paired Student's t-test).

**Figure S4**

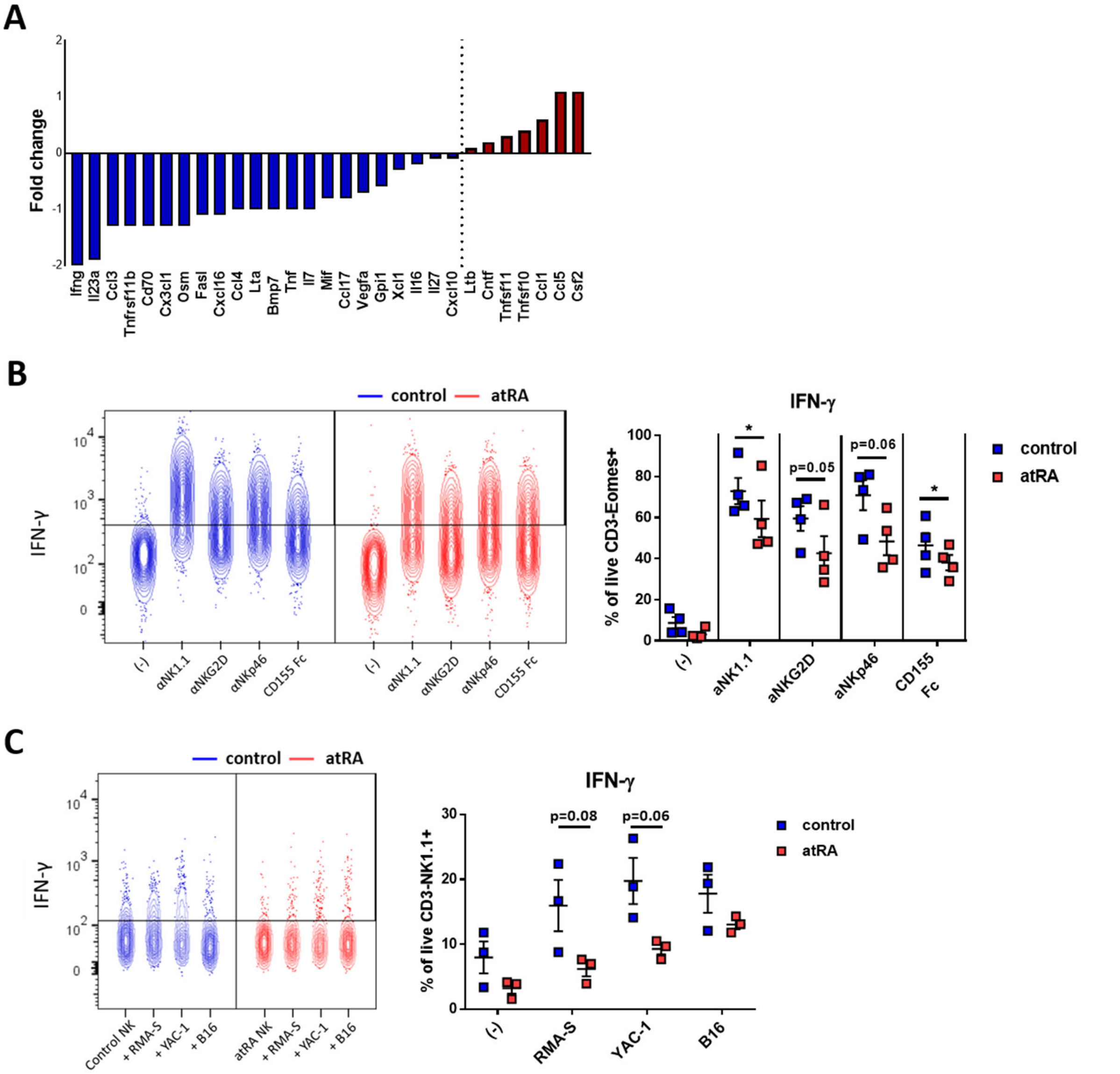


**Figure S4. atRA-treated NK cells produce less of IFN-γ than control NK cells upon stimulation.**

**(A)** NK cells were cultured with 1 μM of atRA (or DMSO as a control solvent) in the presence of IL-2 for 7 days. Relative mRNA expression of indicated cytokine- and chemokine-encoding transcripts by atRA-treated NK cells normalized to expression of control NK cells.
**(B-C)** Representative contour-plots (right) and quantification (left) of IFN-γ-expressing NK cells, in response to activating receptor triggering (n=4) (B), or tumor cells (n=3) (C). The graphs indicate mean ± SEM; *p<0.05, **p<0.01, ***p<0.001 by paired Student's t-test.

**Figure S5**

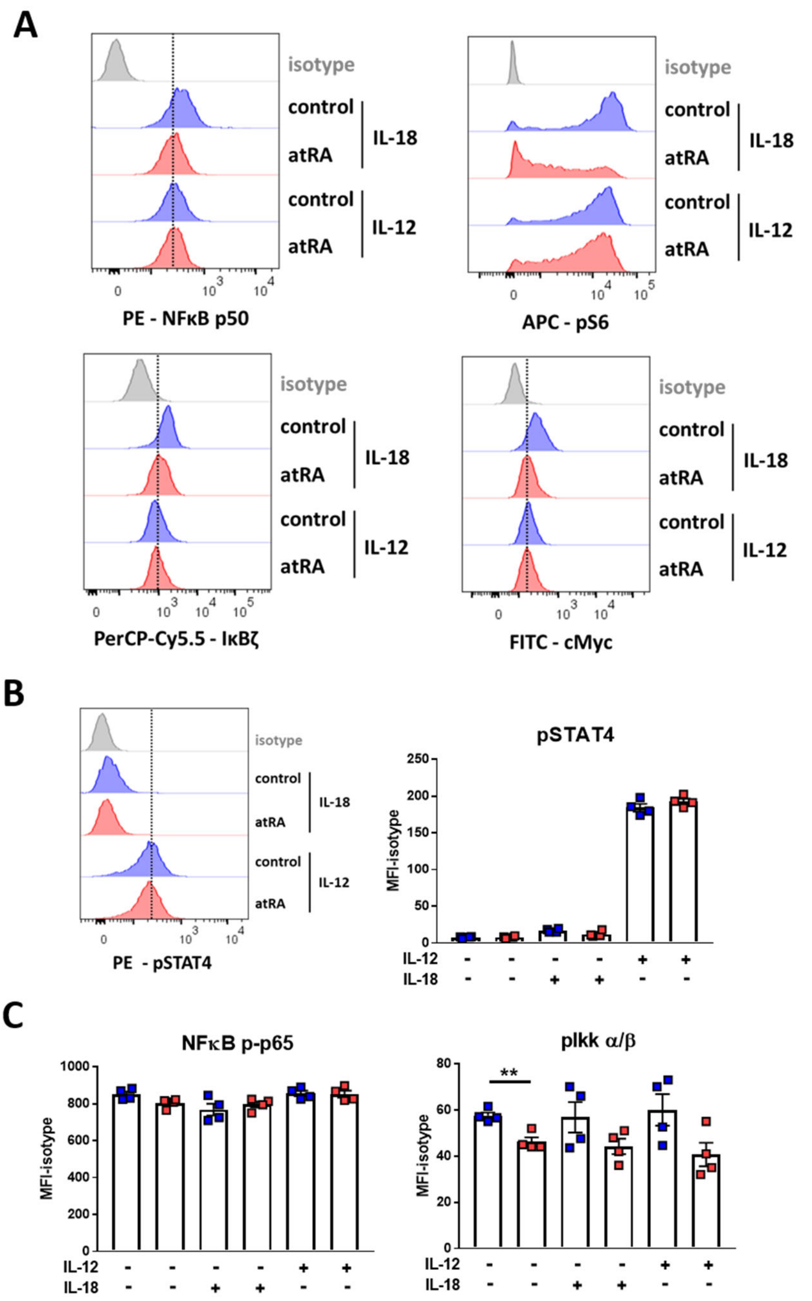


**Figure S5. Activation of IL-12R, IL-18R and mTORC1 pathway, and cMyc expression in atRA-treated and control NK cells upon stimulation with cytokines.**

NK cells were cultured with 1 µM of atRA (or the equivalent volume of DMSO as a control solvent) in the presence of IL-2 for 7 days, and then stimulated with IL-18 or IL-12 for additional 4 hours.

**(A)** Representative histograms showing expression of NFκB p50, IkBζ, phosphorylated (p)-S6 and c-Myc).

**(B-C)** Expression of phosphorylated (p)-STAT4, NFκB p-p65, and p-IKK α/β was analyzed by flow cytometry. Representative histograms (left) and quantification (right) of (A) p-STAT4, and (B) NFκB p-p65 and p-IKK α/β expression by NK cells are shown (n=3-4; mean ± SEM; not significant by paired Student's t-test).

### Figure S6

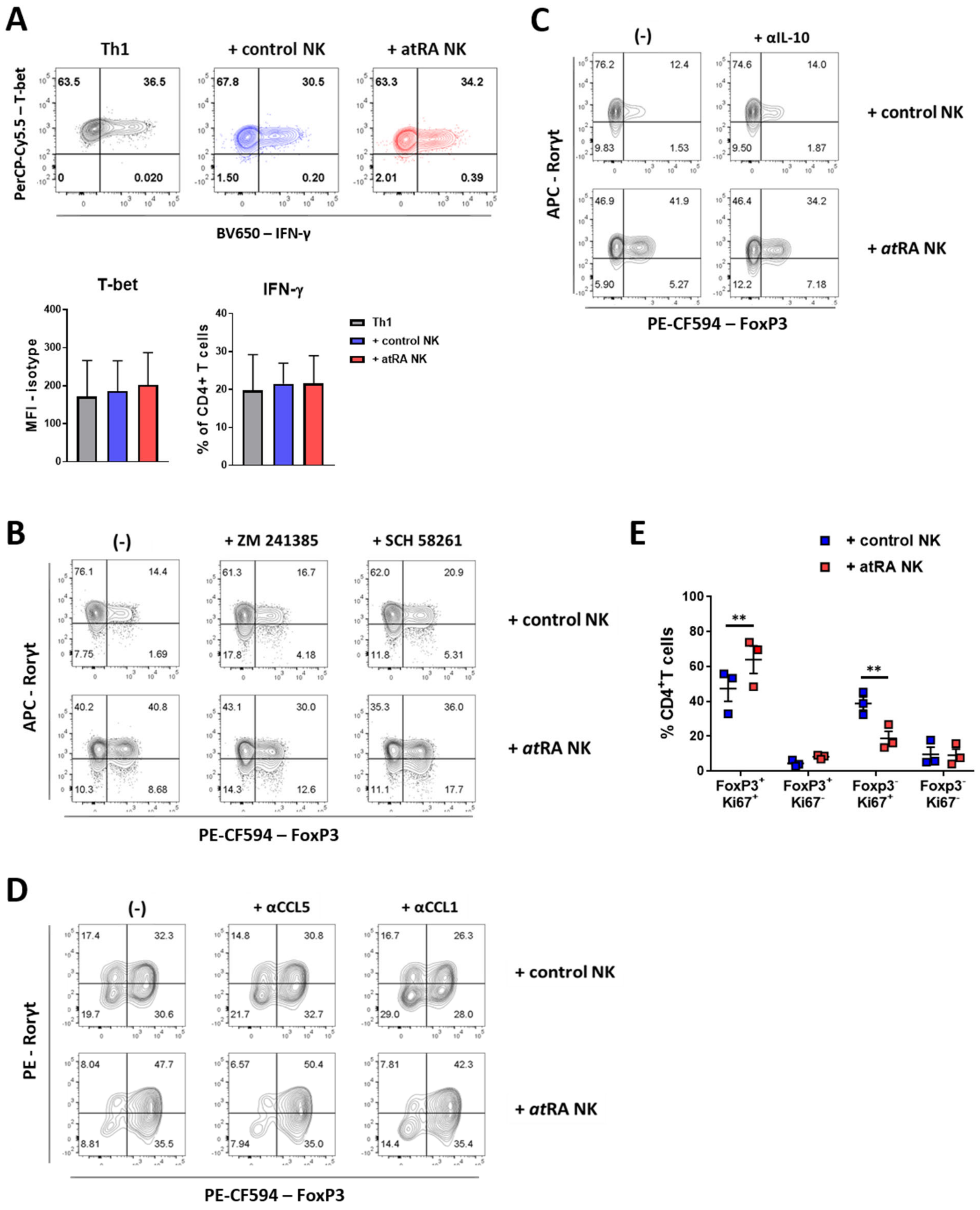


**Figure S6. The co-culture with atRA-treated NK cells does not alter the differentiation of Th1 cells.**

**(A)** CD45.1$^{+}$ NK cells were treated with 1 µM of atRA (or the equivalent volume of DMSO as a control solvent) in the presence of IL-2 for 5 days. NK cells were then co-cultured with naïve CD45.2$^{+}$ CD4$^{+}$ T cells in Th1-polarizing conditions. Upon culture, cells were stimulated with PMA and ionomycin for 4 hours and analyzed by flow-cytometry. Representative contour-plots (up) and quantification (down) of T-bet expression and IFN-γ-production by $CD4^+$ T cells (n=3, mean + SEM; not significant by paired Student's t-test).

**(B-E)** $CD45.1^+$ NK cells were treated with 1 μM of atRA (or the equivalent volume of DMSO as a control solvent) in the presence of IL-2 for 5 days, and then co-cultured with naïve $CD45.2^+$ $CD4^+$ T cells in Treg-polarizing conditions.

**(B-D)** A2AR agonists, ZM 241385 or SCH 58261 (B), anti-IL-10 (C), anti-CCL5 or anti-CCL1 antibodies (D), were added to the co-culture of NK cells and T cells. Contour-plots show Rorγt and FoxP3 expression by $CD4^+$ T cells upon co-culture.

**(E)** Frequencies of Ki67-expressing $FoxP3^+$ or $FoxP3^{neg}$ T cells upon co-culture with NK cells (n=3, mean ± SEM; **, p<0.01 by paired Student's t-test).
